# Scalable Antigen-Antibody Binding Affinity Landscape: A Case Study with ENHERTU

**DOI:** 10.1101/2024.07.12.603351

**Authors:** Wei Li

## Abstract

Optimization of binding affinities for antibody-drug conjugates (ADCs) is inextricably linked to their therapeutic efficacy and specificity, where the majority of ADCs are engineered to achieve equilibrium dissociation constants (K_d_ values) in the range of 10^−9^ to 10^−10^ M. Yet, there is a paucity of published data delineating the optimal binding affinity or its range that ensures improved therapeutic outcomes for ADCs. This study addresses this issue by integrating structural biophysics within a scalable in silico workflow to generate antigen-antibody binding affinity landscapes, with a focus on Trastuzumab, a monoclonal antibody employed in the treatment of HER2-positive breast cancer. By leveraging high-throughput computational techniques, including homology structural modeling and structural biophysics-based K_d_ calculations, this research puts forward a set of high-accuracy structural and intermolecular binding affinity data for Her2-Trastuzumab-Pertuzumab (PDB entry 6OGE). Beyond the design of Her2-targeting ADCs with enhanced efficacy and specificity, this scalable antigen-antibody binding affinity landscape also offers a technically feasible workflow for the high-throughput generation of synthetic structural and biophysical data with reasonable accuracy. Overall, in combination with artificial intelligence (e.g., deep learning) algorithms, this synthetic data approach aims to catalyze a paradigm shift in the discovery and design of antibodies and ADCs with improved efficacy and specificity.

**SIGNIFICANCE:** With Trastuzumab as an example, this study presents a scalable computational biophysical generation of antigen-antibody binding affinity landscapes, serving two purposes: design of Her2-targeting ADCs with enhanced efficacy and specificity and continued accumulation of synthetic structural biophysics data for the development of useful AI-based drug discovery and design model in future. This scalable approach is broadly applicable to databases such as Protein Data Bank.

## INTRODUCTION

Antibody-drug conjugates (ADCs) represent a rapidly advancing class of targeted cancer therapies that merge the specificity of monoclonal antibodies with the potent cytotoxicity of small molecule drugs (1, 2). These biopharmaceuticals are designed to selectively deliver therapeutic agents (i.e., payloads) to cancer cells, thereby minimizing off-target effects and enhancing treatment efficacy (3, 4). To date, drug discovery and design remains a complex multiparameter optimization challenge (5, 6). For instance, central to the efficacy and specificity of ADCs is the binding interaction between the antibody and its target antigen, where the physical strength of this binding is typically quantified as the antigen-antibody equilibrium dissociation constant (K_d_) (7, 8). The K_d_ of ADCs to their target antigens is a critical determinant of receptor-mediated endocytosis and, consequently, the therapeutic efficacy of ADCs (9). ADCs with high binding affinity are more efficiently internalized, as strong binding ensures that the ADC remains attached to the antigen long enough for the endocytic machinery to recognize and internalize the ADC-antigen complex. This leads to more effective delivery of the cytotoxic payload into tumor cells and enhanced specificity, reducing off-target effects and improving the therapeutic index. Conversely, ADCs with low binding affinity may dissociate from the antigen before endocytosis occurs, resulting in reduced internalization and decreased efficacy. Lower affinity might also increase the risk of binding to non-target molecules, leading to off-target effects. Nonetheless, extremely high antigen-antibody affinity can hinder ADC tissue penetration (10), while too low affinity can result in insufficient internalization (11). While most ADCs aim for a K_d_ within the nanomolar range (10^−9^ to 10^−10^ M), few published data have solved the relationship between optimal antigen-antibody K_d_ and the efficacy and the specificity of ADCs (12, 13).

As a result, this study address this paucity of data by developing a scalable in silico approach for structural biophysics-based calculations of antigen-antibody binding affinities, with trastuzumab as an example, a monoclonal antibody widely used in the treatment of HER2-positive breast cancer (14, 15). Experimentally determining for ADCs poses significant challenges, as tools such as surface plasmon resonance and isothermal titration calorimetry necessitate meticulous control of experimental conditions and the use of purified components, making these approaches rather resource- and labor intensive (16–19). On the other hand, computational tools offer a useful alternative approach to explore the antigen-antibody sequence space and chart out the entire territories of antigen-antibody binding affinity landscapes (20). Thus, this study employs computational tools such as structural modeling (21) and physics-based K_d_ calculations (22, 23) to define and build a scalable antigen-antibody binding affinity (K_d_) landscape for the design of antibodies and ADCs with improved efficacy and specificity.

## MATERIALS AND METHODS

Trastuzumab deruxtecan (ENHERTU^®^) is a HER2-directed antibody and DNA topoisomerase I inhibitor conjugate developed for the treatment of HER2-expressing solid tumours (24–26). As the monoclonal antibody in ENHERTU^®^, trastuzumab binds directly to the extracellular domain of the HER2 receptor, inhibiting its downstream signaling pathways and mediating antibody-dependent cellular cytotoxicity (14, 15). As of July 13, 2024, there is a total of 22 trastuzumab-related structures in the Protein Data Bank (PDB) (27, 28), as listed in Table 1.

**Table 1:** Experimentally determined trastuzumab-related structures (released newest from oldest) in PDB as of July 13, 2024, QUERY code: <u>QUERY: Full Text = “Trastuzumab”</u>.

| PDB ID | Structure Title (release date from newest to oldest) |
| --- | --- |
| 8PWH | Atomic structure and conformational variability of the HER2-Trastuzumab-Pertuzumab complex |
| 8Q6J | Atomic structure and conformational variability of the HER2-Trastuzumab-Pertuzumab complex |
| 7PKL | Mechanistic understanding of antibody masking with anti-idiotypic antibody fragments |
| 7MN8 | Structure of the HER2/HER3/NRG1b Heterodimer Extracellular Domain bound to Trastuzumab Fab |
| 6OGE | Cryo-EM structure of Her2 extracellular domain-Trastuzumab Fab-Pertuzumab Fab complex |
| 6BGT | Structure of Trastuzumab Fab mutant in complex with Her2 extracellular domain |
| 6BAH | Trastuzumab Fab v3 with 5-diphenyl mediotope variant |
| 6BAE | Trastuzumab Fab v3 in complex with CQFDLSTRRLKC |
| 6B9Z | Trastuzumab Fab v3 |
| 6B9Y | Trastuzumab Fab v3 in complex with 5-phenyl mediotope variant |
| 5U6A | Crystal structure of I83E mediotope-enabled trastuzumab with azido-peg3-meditope |
| 5U5M | Crystal structure of I83E mediotope-enabled trastuzumab with azido-meditope |
| 5U5F | Meditope enabled trastuzumab I83E variant in complex with Ac) CQFDA(PH)2STRRLRCGGSK |
| 5U3D | Structure of mediotope enabled trastuzumab I83E variant |
| 6BI2 | Trastuzumab Fab D185A (Light Chain) Mutant Biotin Conjugation |
| 6BI0 | Trastuzumab Fab N158A, D185A, K190A (Light Chain) Triple Mutant |
| 6BHZ | Trastuzumab Fab D185A (Light Chain) Mutant |
| 5XHG | Crystal structure of Trastuzumab Fab fragment bearing Ne-(o-azidobenzyloxycarbonyl)-L-lysine |
| 5XHF | Crystal structure of Trastuzumab Fab fragment bearing p-azido-L-phenylalanine |
| 4IOI | Meditope-enabled trastuzumab in complex with CQFDLSTRRLKC |
| 4HKZ | Trastuzumab Fab complexed with Protein L and Protein A fragments |
| 4HJG | Meditope-enabled trastuzumab |

Among the 22, there are a total of three HER2-Trastuzumab-Pertuzumab complex structures with PDB IDs <u>6OGE</u> (14)), 8PWH (15) and 8Q6J (15). In light of the standardized PDB data format for biomolecular structures, it does not really matter which one of the three is chosen here for subsequent structural modeling (21) and physics-based K_d_ calculations (22, 23), as all three HER2-Trastuzumab-Pertuzumab complex structures are determined experimentally with Cryo-EM (14, 15). Moreover, while only trastuzumab is used in ENHERTU^®^, a synergistic anticancer effect of the two antibodies is also likely, according to a detailed CryoEM study of the ternary complex (15). Thus, this study here chooses PDB entry <u>6OGE</u> (14)) as an example to define and build a scalable antigen-antibody binding affinity (K_d_) landscape. Briefly, the relationships between chain IDs and molecular entities of PDB entry 6OGE are listed in Table 2, which is to be used in describing the results of the subsequent structural modeling (21) and physics-based K_d_ calculations (22, 23).

**Table 2:** Relationships between chain IDs and molecular entities of PDB entry 6OGE. In this table, length represents the number of amino acid residues, total means A+B+C+D+E.

| Chain ID | Molecular entity | Length |
| --- | --- | --- |
| A | Receptor tyrosine-protein kinase erbB-2 (Homo sapiens, 9606) | 622 |
| B | Pertuzumab FAB LIGHT CHAIN (Homo sapiens, 9606) | 214 |
| C | Pertuzumab FAB HEAVY CHAIN (Homo sapiens, 9606) | 222 |
| D | Trastuzumab FAB LIGHT CHAIN (Homo sapiens, 9606) | 214 |
| E | Trastuzumab FAB HEAVY CHAIN (Homo sapiens, 9606) | 220 |
| BC | Pertuzumab FAB (Homo sapiens, 9606) | 436 |
| DE | Trastuzumab FAB (Homo sapiens, 9606) | 434 |
| Total | PDB entry 6OGE (14) | 1492 |

With PDB entry 6OGE (14) (Table 1) as an initial input, subsequent structural modeling (21) and physics-based K_d_ calculations (22, 23) consists of an automated in silico generation of synthetic homology structural and K_d_ data, as illustrated in Figure 1 and described previously in detail (29). Briefly, Modeller (21) was employed to build a total of 29,840 (1, 492 × 20) homology structural models one site-specific missense mutation introduced to PDB entry 6OGE (14). Afterwards, the binding affinities were calculated using Prodigy (22, 23) for all 29,840 structural models of Her2-Trastuzumab-Pertuzumab analogues, including the K_d_ values between chains A and B, chains A and C, chains A and D, chains A and E, chains A and BC, chains A and DE (Table 2). With PDB entry 6OGE (14) as template, all structural modeling (21) and physics-based K_d_ calculations (22, 23) were repeated three times on Wuxi Taihu Lake High Performance Computing platforms.

**Figure 1:**
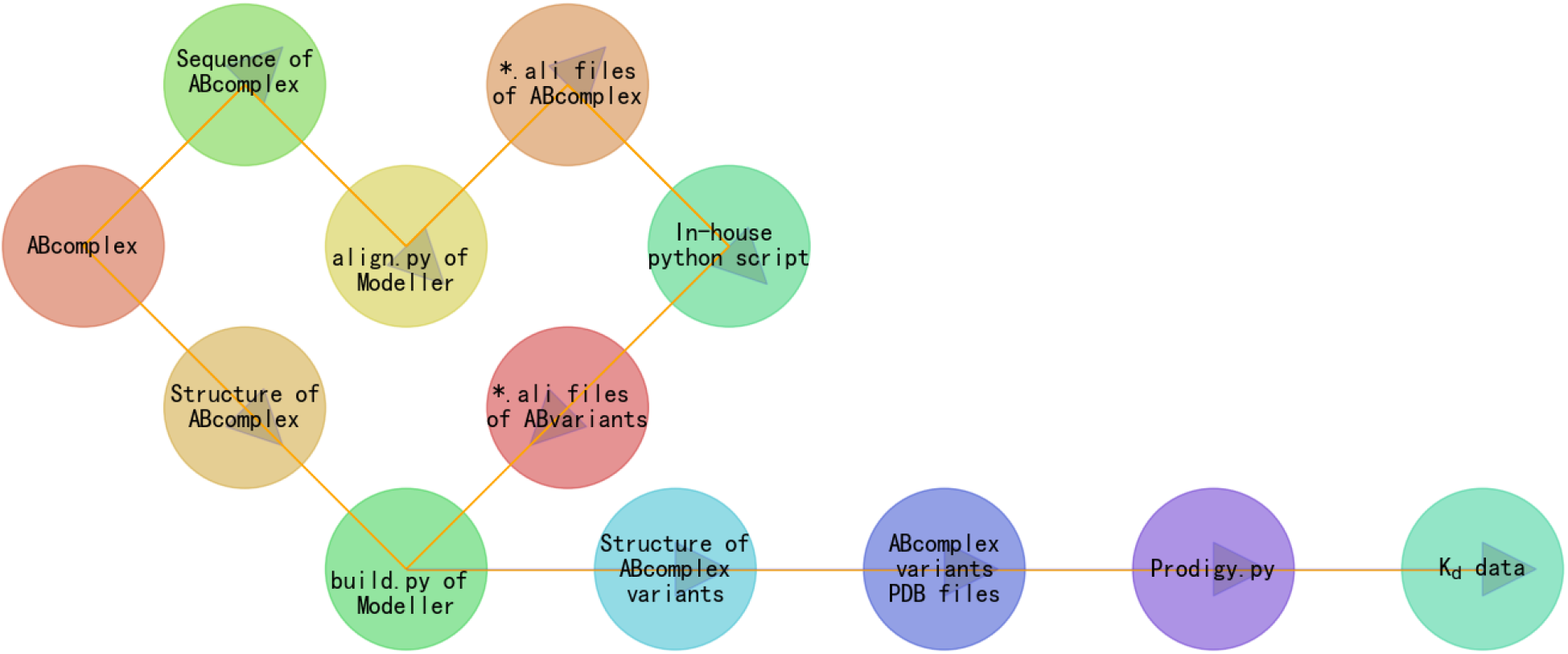
Automated in silico generation of synthetic structural (Modeller) and K_d_ (Prodigy) data.

Of further note, among the twenty natural amino acids, cysteine is a special one from a structural point of view, in the sense that removal of cysteine residue(s) or introduction of new cysteine residue(s) might induce a perturbation of the disulfide bonding network towards a major structural rearrangement of a protein. Yet, engineering cysteines at specific sites in antibodies has proven a promising approach to create well-defined ADCs for the treatment of cancer (30, 31). This study, therefore, reports a computational systematic amino acid (including cysteine) scanning (13, 32–35) of the entire PDB entry 6OGE (1492 amino acid residues, Table 2) (14) to incorporate structural biophysics (e.g., K_d_) (8) into the property-based design of cysteine-linked ADCs (5, 6).

## RESULTS

Since this study used PDB entry 6OGE (14) as the structural template, Prodigy (22, 23) was used to calculate the inter-chain binding affinities for the native experimental Her2-Trastuzumab-Pertuzumab complex structure. These physics-based calculations were performed between chains A and B, chains A and C, chains A and D, chains A and E, chains A and BC, and chains A and DE, as listed in Table 3 as below.

**Table 3:** Inter-chain binding affinities calculated by Prodigy for the native experimental Her2-Trastuzumab-Pertuzumab complex structure PDB entry 6OGE. In this table, the codes (A, B, C, etc) for binding partners are defined in Table 2.

| Binding partners | Inter-chain binding affinity (M) at 37 °C |
| --- | --- |
| A + B | $1.4 \times 10^{-5}$ |
| A + C | $9.5 \times 10^{-8}$ |
| A + D | $8.6 \times 10^{-6}$ |
| A + E | $3.8 \times 10^{-6}$ |
| A + BC | $1.9 \times 10^{-8}$ |
| A + DE | $4.9 \times 10^{-7}$ |

Afterwards, with PDB entry 6OGE (14) as the structural template, this study conducted three sets of structural modeling (21) and physics-based K_d_ calculations (22, 23). These calculations were also performed to determine K_d_ values for interactions between chains A and B, chains A and C, chains A and D, chains A and E, chains A and BC, and chains A and DE, as detailed in Table 2. The resulting K_d_ values and their corresponding analyses are comprehensively summarized in Table 4, with additional data presented in six supplementary tables and six figures contained in the supplementary file <u>supps.pdf</u>.

**Table 4:** Scalable Her2-Trastuzumab-Pertuzumab binding affinity landscapes and their distribution patterns. In this table, supplementary file represents <u>supps.pdf</u>, the codes (A, B, C, etc) for binding partners are defined in Table 2.

| Molecule pair | Scalable $K_d$ landscape | $K_d$ distribution histogram |
| --- | --- | --- |
| A + B | Table 3 (supplementary file) | Figure 1 (supplementary file) |
| A + C | Table 4 (supplementary file) | Figure 2 (supplementary file) |
| A + D | Table 5 (supplementary file) | Figure 3 (supplementary file) |
| A + E | Table 6 (supplementary file) | Figure 4 (supplementary file) |
| A + BC | Table 7 (supplementary file) | Figure 5 (supplementary file) |
| A + DE | Table 8 (supplementary file) | Figure 6 (supplementary file) |

According to Table 3, the K_d_ between Her2 and pertuzumab Fab is located at 1.9 × 10^−8^ M (vertical red line in Figure 2) for the native complex structure of Her2-Trastuzumab-Pertuzumab (PDB entry 6OGE), while the K_d_ values between Her2 and pertuzumab Fab possesses a much wider distribution, ranging from 1.9 × 10^−7^ M to 2.9 × 10^−10^ M, according to the Prodigy (22, 23) calculations of the 3 × 29,840 homology structural models of the Her2-Trastuzumab-Pertuzumab complex (PDB entry 6OGE) with one site-specific missense mutation.

**Figure 2:**
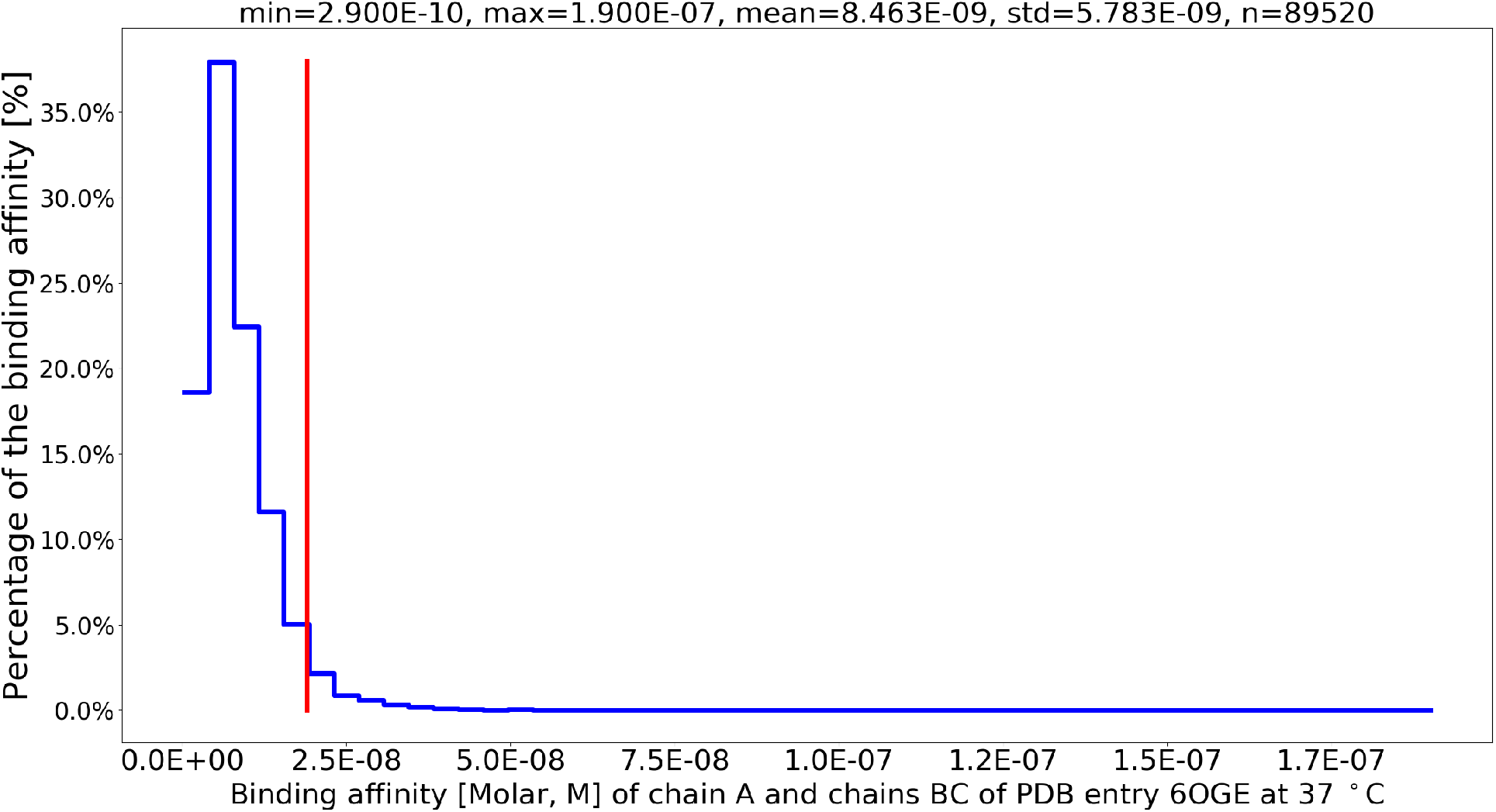
Histogram depicting the distribution pattern of the Her2-Pertuzumab binding affinities between chain A and chain BC (listed in Table 2) of PDB entry 6OGE with one site-specific missense mutation. The vertical red line indicates the K_d_ between chain A (Her2) and chain BC (Pertuzumab FAB), representing the native complex structure of Her2-Trastuzumab-Pertuzumab (PDB entry 6OGE).

Similarly, according to Table 3, the K_d_ between Her2 and trastuzumab Fab is located at 4.9 × 10^−7^ M (vertical red line in Figure 3) for the native complex structure of Her2-Trastuzumab-Pertuzumab (PDB entry 6OGE), while the K_d_ values between Her2 and trastuzumab Fab also possesses a much wider distribution, ranging from 2.5 × 10^−6^ M to 1.7 × 10^−8^ M, according to the Prodigy (22, 23) calculations of the 3 × 29,840 homology structural models of the Her2-Trastuzumab-Pertuzumab complex (PDB entry 6OGE) with one site-specific missense mutation.

**Figure 3:**
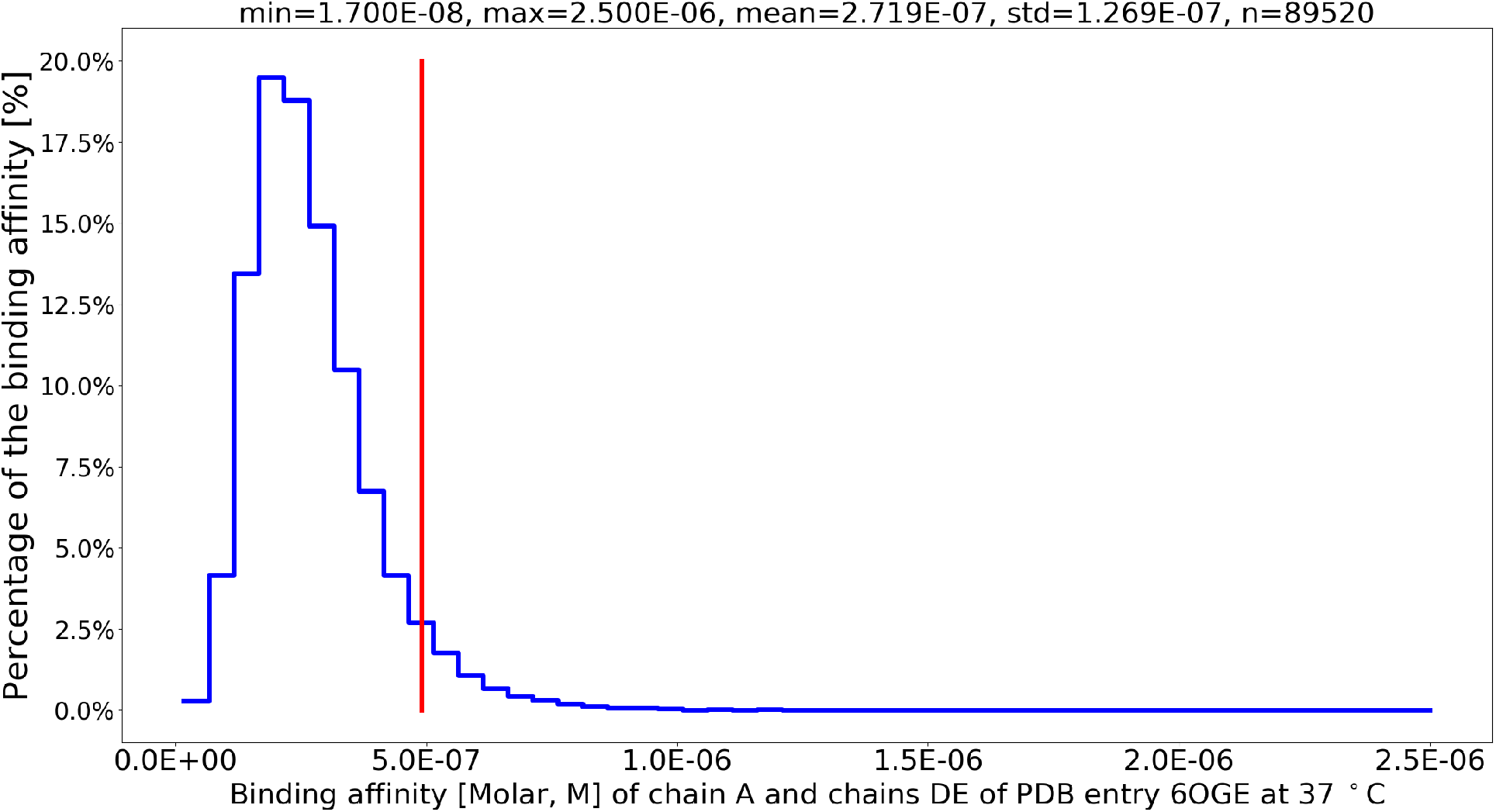
Histogram showing the distribution pattern of the Her2-Trastuzumab binding affinities between chain A and chain DE (listed in Table 2) of PDB entry 6OGE with one site-specific missense mutation. The vertical red line marks the K_d_ between chain A (Her2) and chain DE (Trastuzumab FAB), representing the native complex structure of Her2-Trastuzumab-Pertuzumab (PDB entry 6OGE).

Taken together, for both pertuzumab Fab and trastuzumab Fab, there are rooms for both increase and decrease of their antigen-antibody K_d_ values, as shown in Figures 2 and 3, and the two scalable antigen-antibody binding affinity landscapes (Figures 2 and 3) are like two binding affinity maps, which is one inextricable factor in the equation of *y* = *f* (*x*), where *x* stands for the optimal K_d_ or a range of it, while *y* stands for optimal efficacy and specificity of antibodies and ADCs (12, 13).

Of an interesting note, the HER2-Trastuzumab-Pertuzumab binding affinity landscapes (Figures 2 and 3) elucidate the binding affinity variations caused by site-specific mutations, such as the <u>S911F</u> mutation in chain C of the pertuzumab heavy chain, as demonstrated in the supplementary file <u>fin.pdb</u> and Figure 7 of supplementary file <u>supps.pdf</u>. After the computational analysis on Wuxi Taihu Lake High Performance Computing platforms, this particular mutation <u>S911F</u> was also assessed using the Prodigy server (22, 23), which returned an identical K_d_ between chain A (Her2) and chain BC (Pertuzumab FAB, Table 2) of 2.9 × 10^−10^ M, mutation No. 18214 in Table 7 of supplementary file <u>supps.pdf</u>. This K_d_ of 2.9 × 10^−10^ M indicates an ∼ two orders of magnitude enhanced antigen-antibody binding affinity due to this particular mutation <u>S911F</u>, compared to the K_d_ of 1.9 × 10^−8^ M between Her2 and pertuzumab Fab in the native experimental Her2-Trastuzumab-Pertuzumab complex structure, according to Table 3.

## DISCUSSION

This study employs a previously described scalable in silico workflow (29), including structural modeling (21) and physics-based K_d_ calculations (22, 23), to define and build two scalable antigen-antibody binding affinity landscapes (Figures 2 and 3) with Her2-Trastuzumab-Pertuzumab as an example (PDB entry 6OGE (14)). This scalable in silico workflow (29) provides a novel perspective for the application of computational structural biophysics in therapeutic antibody and ADC design, because the introduction of only one site-specific missense mutation ensures reasonable accuracy of the two Her2-Trastuzumab-Pertuzumab binding affinity landscape (Figures 2 and 3), which consists of a huge set of structural and biophysical calculations (21–23).

Of further note, this Her2-Trastuzumab-Pertuzumab binding affinity landscape (Figures 2, 3 and supplementary information in Table 4) is also able to be used the other way around, i.e., to be used as a search engine (8) for K_d_-ranked Trastuzumab or Pertuzumab analogues in the discovery and design of next-generation Her2-targeting ADCs (12, 13). Moreover, this scalable in silico workflow (29) also offers a technically feasible approach for high-throughput generation of synthetic structural and biophysical data with reasonable accuracy (8, 36). To this end, it is conceivable that this synthetic data approach, in combination with artificial intelligence (AI, e.g., deep learning) algorithms, is able to catalyze an AI-accelerated paradigm shift in the discovery and design of therapeutic antibodies and ADCs with improved efficacy and specificity (5, 6, 37).

In short, this study starts from an experimental Her2-Trastuzumab-Pertuzumab complex structure (PDB entry 6OGE (14)) to build two scalable antigen-antibody binding affinity landscapes (Figures 2 and 3), which are scalable because:

1. this Modigy (Figure 4) workflow (29) is broadly applicable to biomolecular strcutre databases such as PDB (27) and AFDB (38–41).
2. this Modigy (Figure 4) workflow (29) introduced only <u>one</u> site-specific missense mutation to the Her2-Trastuzumab-Pertuzumab complex structure (PDB entry 6OGE (14)), where the number could be larger, as long as the overall accuracy is reasonable for the synthetic structural and biophysical data (29).
3. the Her2-Trastuzumab-Pertuzumab binding affinity landscape (Figures 2 and 3) includes not only site-specific mutants of the two antibodies, but also site-specific mutants of the target, i.e., Her2 as the antigen, highlighting the use of this in silico workflow (29) in high-throughput generation of synthetic structural and biophysical data for other drug targets (GPCRs (42), ion channels (43), etc.) to train AI models for the discovery and design (8) of not just therapeutics antibodies and ADC, but also of small molecule compounds (44, 45).
4. method-wise, in addition to the structural modeling (21) and physics-based K_d_ calculations (22, 23) employed here, molecular dynamics simulations (46, 47) is also rather useful to further enhance the accuracy of the scalable antigen-antibody binding affinity landscape for drug discovery and design in future (48, 49).

**Figure 4:**
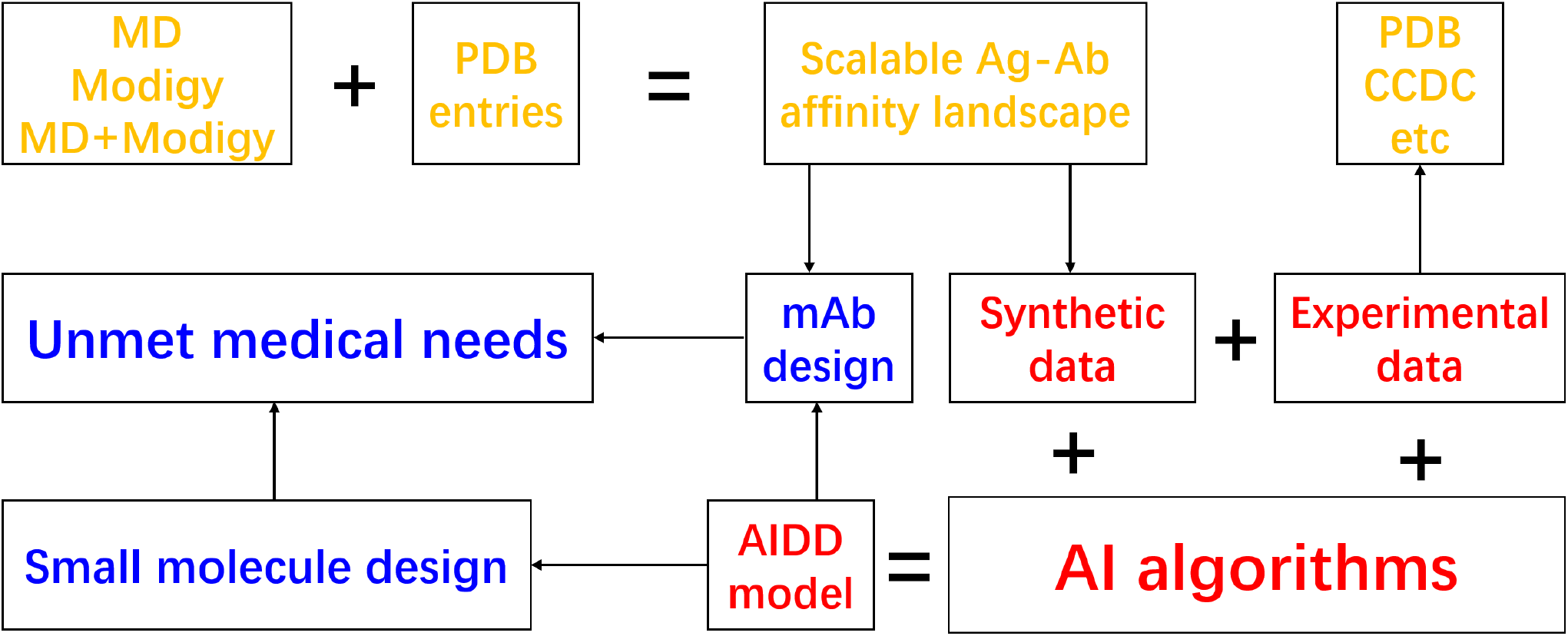
A flowchart depicting the generation of a scalable antigen-antibody binding affinity landscape for designing antibodies and small molecule compounds with improved efficacy and specificity. In this figure, Modigy represents the method approach combining Modeller and Prodigy for in silico generation of structural and intermolecular binding affinity data, as described in the Methods section.

To sum up, the synthetic structural and biophysics data serve two purposes: (1), this scalable Modigy (Figure 4) workflow (29) creates a scalable antigen-antibody binding affinity landscape, which acts like a map to guide the design of monoclonal antibodies or ADCs with optimal binding affinities (12, 13); (2), this scalable Modigy (Figure 4) workflow (29) generates useful training data (50, 51) for AI-driven drug design (AIDD, Figure 4) models (8). Finally, while this scalable Modigy (Figure 4) workflow (29) facilitates the rational design of next-generation therapeutic antibodies and ADCs with enhanced efficacy, specificity, future AIDD models are also to further enhance the design of both monoclonal antibodies, ADCs (12, 13) and also small molecule compounds (Figure 4) with improved efficacy and specificity to better address unmet medical needs (5, 6).

## CONCLUSION

To sum up, with ENHERTU^®^’s Trastuzumab as an example, this study presents a scalable computational biophysical generation of antigen-antibody binding affinity landscapes, serving two purposes: design of Her2-targeting ADCs with enhanced efficacy and specificity and continued accumulation of synthetic structural biophysics data for the development of useful AI-based drug discovery and design model in future. This scalable approach is broadly applicable to databases such as Protein Data Bank.

## Supporting information

supplementary tables and figures of the antigen-antibody binding affinity landscapes

## AUTHOR CONTRIBUTIONS

Conceptualization, W.L.; methodology, W.L.; software, W.L.; validation, W.L.; formal analysis, W.L.; investigation, W.L.; resources, W.L.; data duration, W.L.; writing–original draft preparation, W.L.; writing–review and editing, W.L.; visualization, W.L.; supervision, W.L.; project administration, W.L.;

## ACKNOWLEDGMENTS

The author is grateful to the communities of structural biology, biophysics, medicinal and computational chemistry and algorithm design, for the continued accumulation of knowledge and data for drug discovery & design, and for the continued development of tools (hardware, software and algorithm) for drug discovery & design.

## SUPPLEMENTARY MATERIAL

1. Supplementary file <u>fin.pdb</u>: a homology structural model of the Her2-Trastuzumab-Pertuzumab complex (PDB entry 6OGE) with one particular mutation <u>S911F</u>.
2. Supplementary file <u>supps.pdf</u>: supplementary tables and figures of the antigen-antibody binding affinity landscapes with ENHERTU^®^’s Trastuzumab as an example.

